# Deep sequencing reveals *Campylobacter* in commercial meat chickens less than 8 days old

**DOI:** 10.1101/2020.06.10.141978

**Authors:** F.M. Colles, S.J. Hedges, R. Dixon, S.G. Preston, P. Thornhill, K.K. Barfod, S.G. Gebhardt-Henrich, P. Creach, M.C.J. Maiden, M.S. Dawkins, A.L. Smith

## Abstract

*Campylobacter* from contaminated poultry meat is a major source of human gastroenteritis worldwide. To date, attempts to control this zoonotic infection with on-farm biosecurity measures have been inconsistent. A cornerstone of these efforts has been the detection of chicken infection with microbiological culture, which typically does not occur until birds are at least 21 days old. Using molecular methods for detecting *Campylobacter* presence, 16S microbiome analysis and deep sequencing of the *Campylobacter por*A gene, *Campylobacter* can be identified at very low levels in most or all flocks fewer than 8 days old. These young chicks exhibit a much greater diversity of *porA* types than older birds testing positive for *Campylobacter* by culture or qPCR. This suggests that, as the bacteria multiply sufficiently to be detected by culture methods, one or two strains, as indicated by *porA* type, dominate the infection. The findings that (i) most young chicks carry some *Campylobacter* and (ii) not all flocks become *Campylobacter* positive by culture, suggests that efforts to control infection should concentrate on how to maintain *Campylobacter* at low levels by the prevention of the overgrowth of single strains, which ultimately leads to the contamination of food.

## Main

The consumption of raw or undercooked poultry meat contaminated with the bacterium *Campylobacter* is one of the major causes of gastroenteritis within human populations in high-income countries ^1,2^. Current strategies for reducing infection in poultry have primarily focussed on increasing biosecurity on-farm and preventing the infection of chicken flocks by cross-contamination. These approaches have been inconsistent as an intervention and levels of infection have remained high at around 70% of EU flocks being contaminated^2,3^. Detecting the presence of *Campylobacter* within a flock currently relies on growing the bacteria in culture or by quantitative PCR (qPCR), which generally identify *Campylobacter* positive flocks when the chickens are 3-5 weeks old^4^. These findings led to the widespread assumption that chickens are free from infection before this age, but it is not known whether they are genuinely *Campylobacter*-free or rather that *Campylobacter* are present but not detected because of lack of sensitivity of the methods used. To address this question, we applied different methods for detecting *Campylobacte*r in chickens: (i) microbiological culture or qPCR; (ii) 16S microbiome analysis; and (iii) deep sequencing of the *Campylobacter porA* gene^5^. The culture methods used standard growth media, but the 16S microbiome analysis was conducted with high sequencing depth (average 65,000 sequences/sample) so, that it could detect *Campylobacter* in samples even in small quantities. The *porA* gene was chosen because it encodes for a surface-exposed major outer membrane protein (MOMP) in *Campylobacter* and has been successfully included in typing schemes to assess genetic diversity^6^. Deep sequencing of a fragment of the short variable region of the gene (*porAf2*) was used since this efficiently identifies multiple strains within samples^7^. The 34 commercial broiler (meat) flocks sampled for this study were from the UK, France, and Switzerland representing a range of different housing and management conditions.

## Results

*Campylobacter* DNA was detected in faecal samples from birds less than 8 days of age in all 34 commercial broiler flocks tested across three different European countries, 5 different companies and 17 different farms. Testing up to 5 faecal samples/flock, 87.5% of flocks (39/78 faecal samples) tested positive for *Campylobacter* using high depth 16S rRNA microbiome analysis, and 100% of flocks (58/78 faecal samples) tested positive using the *porAf2* deep sequencing target (Table 2; Figure 1a and b). Between 0 and 6 *Campylobacter* sequence variants were recovered in samples from young birds, with an average sequencing depth of 65,000 per sample (supplementary data, Figure 1). *Campylobacter* DNA was also detected in samples processed individually, controlling for cross-contamination. In contrast, *Campylobacter* DNA was not detected amongst non-template negative controls using the same techniques, or from samples tested from 5 high-biosecurity specific pathogen free chickens tested at 3 or 7 days of age, using an average 16S microbiome sequencing depth of 26,575 reads per sample. Although at the lower end of the sequencing depth achieved for the commercial birds, *Campylobacter* DNA was detected in 13/78 samples in commercial birds at an equivalent sampling depth.

**Table 1.**
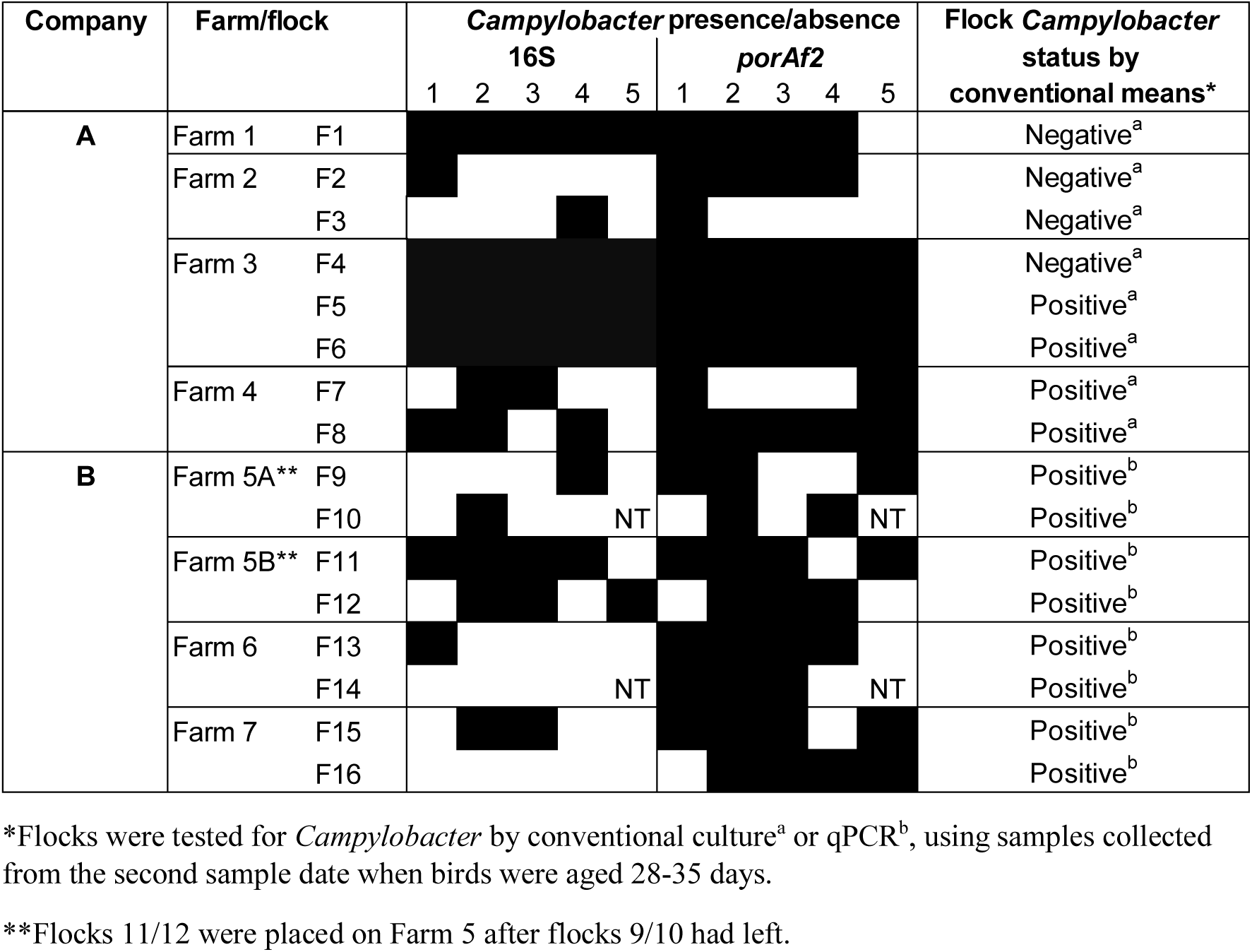
Presence/absence of *Campylobacter* DNA amongst faecal samples from UK broiler chicks < 7 days of age, detected by deep sequencing of the 16S rRNA or *porAf2* targets. Fresh faecal samples were collected from up to 5 birds selected at random at each time point. The 16S rRNA and *porAf2* targets were amplified from DNA extracted directly from individual samples by PCR, and the number of variants detected by deep sequencing. 16S rRNA analysis at high depth was used to look for presence of *Campylobacter* amongst the total microbiome, whilst amplification of the *porAf2* target was used to enrich for *Campylobacter* specific DNA relative to the other microbiota.

**Table 2.** Detection of *Campylobacter* DNA, defined as the percentage of the total number of 16s rRNA microbiome sequences recovered from pooled faecal samples from each of the flocks. The figures shown are the average of samples from 4-5 birds, with standard deviation shown in brackets. Results are shown for a subset of 16 UK flocks from 2 different companies that were tested at early and late time points during rearing. The results are split by flocks that were culture negative and culture positive for *Campylobacter* at the later sample point.

| Flock | *Campylobacter* negative |  | *Campylobacter* positive |  |  |
| --- | --- | --- | --- | --- | --- |
|  | 6-7 days | 34-35 days | 6-7 days | 28 days | 34-35 days |
| *Company A* |  |  |  |  |  |
| Flock 1 | 0.0022 (0.002) | 0.0012 (0.002) |  |  |  |
| Flock 2 | 0.0017 (0.004) | 0.0000 (0.000) |  |  |  |
| Flock 3 | 0.0003 (0.001) | 0.0016 (0.002) |  |  |  |
| Flock 4 | 0.0081 (0.003) | 0.0057 (0.004) |  |  |  |
| Flock 5 |  |  | 0.0120 (0.006) |  | 0.6856 (1.305) |
| Flock 6 |  |  | 0.0099 (0.010) |  | 0.1563 (0.162) |
| Flock 7 |  |  | 0.0005 (0.001) |  | 0.0012 (0.002) |
| Flock 8 |  |  | 0.0007 (0.001) |  | 0.0185 (0.028) |
| *Company B* |  |  |  |  |  |
| Flock 9 |  |  | 0.0005 (0.001) | 0.0008 (0.001) |  |
| Flock 10 |  |  | 0.0001 (0.000) | 0.0002 (0.000) |  |
| Flock 11 |  |  | 0.0014 (0.001) |  | 0.0060 (0.008) |
| Flock 12 |  |  | 0.0018 (0.001) |  | 0.0507 (0.062) |
| Flock 13 |  |  | 0.0003 (0.002) | 0.0011 (0.001) |  |
| Flock 14 |  |  | 0.0000 (0.000) | 0.0008 (0.002) |  |
| Flock 15 |  |  | 0.0012 (0.002) | 0.0063 (0.012) |  |
| Flock 16 |  |  | 0.0000 (0.000) | 0.0146 (0.021) |  |
| *Overall average* | 0.0031 | 0.0021 | 0.0024 | 0.0036 | 0.1377 |

**Figure 1.**
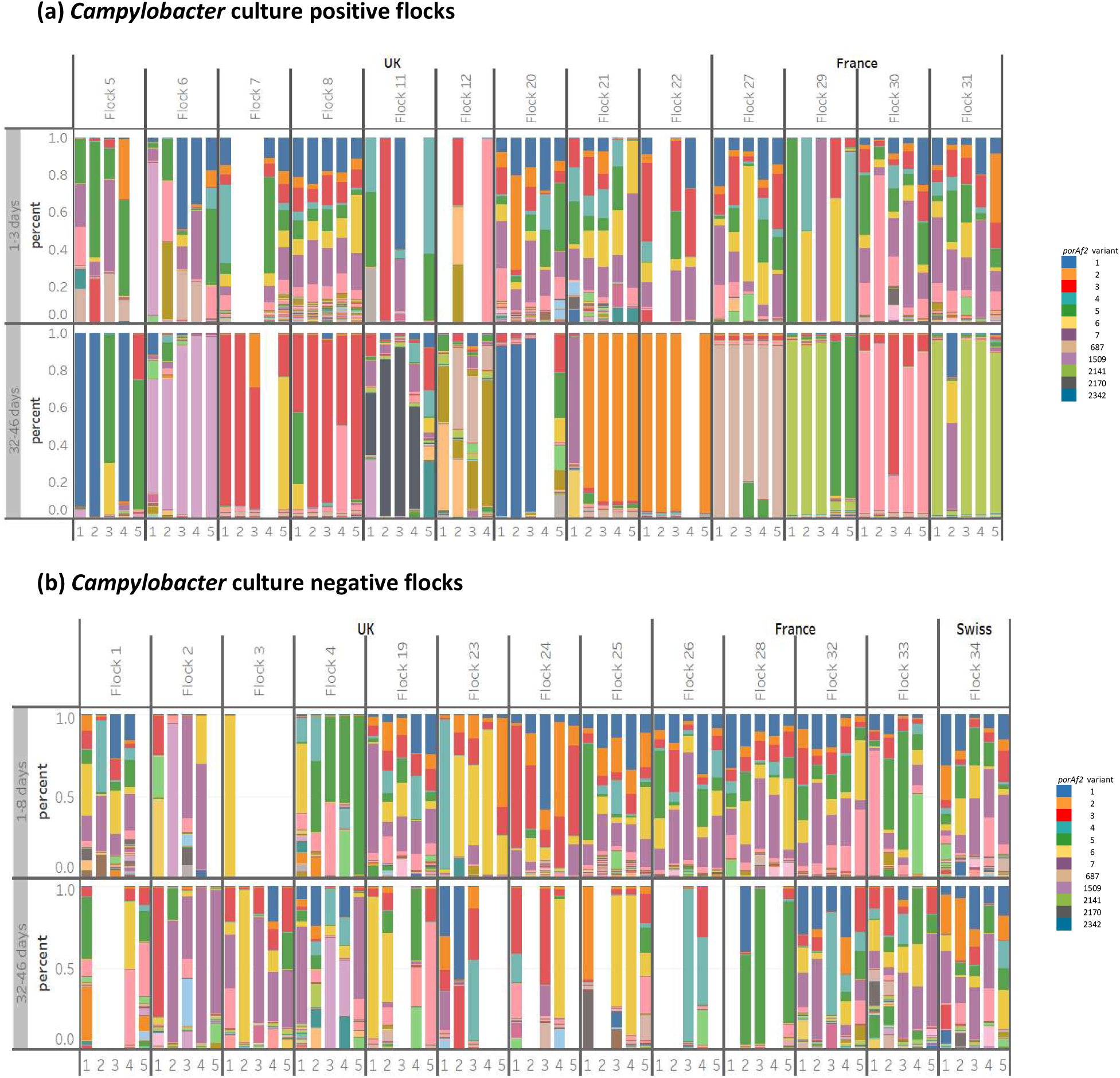
The prevalence of *Campylobacter porAf2* types amongst faecal samples collected from 34 commercially reared broiler flocks from the UK, France, and Switzerland (normalised data). Results are shown for flocks that tested (a) *Campylobacter* positive and (b) *Campylobacter* negative before slaughter by culture/qPCR. Flocks that were tested at 28-31 days of age are not shown, together with flock 14 that had low sequencing depth. Each colour represents a different *porAf2* type; common variants are given in the legend.

The quantities of *Campylobacter* DNA detected in the young commercial bird samples was extremely small, ranging from 0.0001-0.012% of the total sequencing reads (Table 2) and was similar, regardless of whether or not the flocks subsequently tested *Campylobacter* culture or qPCR positive at 28-35 days. However, if a flock tested *Campylobacter* culture or qPCR positive at 35 days, the proportion of *Campylobacter* sequences in the microbiome was 100 fold larger at 0.1377% of the sequencing reads (±0.490 standard deviation) (Wilcoxon rank-sum exact test, N=6 flocks *p* = 0.03). The large deviation reflected a small subset of samples (5/151 samples; 3.3%) from 35 day-old chickens across 3 flocks having >0.1% *Campylobacter* abundance. The proportion of *Campylobacter* sequences/total 16S did not differ significantly between young and older birds from the same flock from *Campylobacter* culture negative flocks (Wilcoxon rank-sum exact text, N=4 flocks, *p* = 0.375).

Despite the low level of *Campylobacter* in young birds (0.0001-0.012% of the microbiome), the *porAf2* sequences were highly variable. A total of 1098 *porAf2* types (1066 in samples from 25 UK flocks; 353 from 8 French flocks; 560 from 1 Swiss flock), corresponding to 746 peptide sequences, were identified amongst the flocks tested in the study (total sequencing data). Rarefaction curves demonstrated that sequencing depth was sufficient to capture the *porAf2* variant diversity from most flocks at young and older points of sampling (supplementary data, Figure 2). Of the 1098 *porAf2* types, 439 (40.0%) were unique to UK flocks and 32 (2.9%) were unique to French flocks. None of the *porAf2* types were unique to the Swiss flock, which is perhaps unsurprising since only one Swiss flock was tested. All *porAf2* types from the Swiss flocks were also identified amongst UK flocks, and 254 were identified French flocks. Seven of 1098 (0.64%) *porAf2* types (*porAf2* types 1-7) were identified in all 34 flocks, accounting for 49.36% (2,315,725/4,691,464) of the total sequences recovered across the whole study (supplementary data, Figures S3 and S4). The number of different *porAf2* types recovered from an individual faecal sample ranged widely from 9 to 358. There was no significant difference in the average number of *porAf2* types recovered from young broiler flocks that tested *Campylobacter* culture-negative at 28-46 days compared to flocks that tested *Campylobacter* positive by standard methods (normalised data, N=33 flocks, Mann-Whitney U test, *p* = 0.6713), or between flocks tested at early and later time points (normalised data, Wilcoxon rank-sum exact test, N=33 flocks, *p* = 0.618) (supplementary data, Figure 5). For *porAf2* types that became predominant within flocks, 76.1% (16/21) were identified amongst younger birds from the flock, and most often (11/21, 52.4%) at higher proportion of the total *porAf2* sequenced compared other young birds from other flocks (Wilcoxon rank-sum exact test, N=13 flocks, *p* = 0.001 within countries and *p =* 0.002 between countries (Table 4).

**Table 3.**
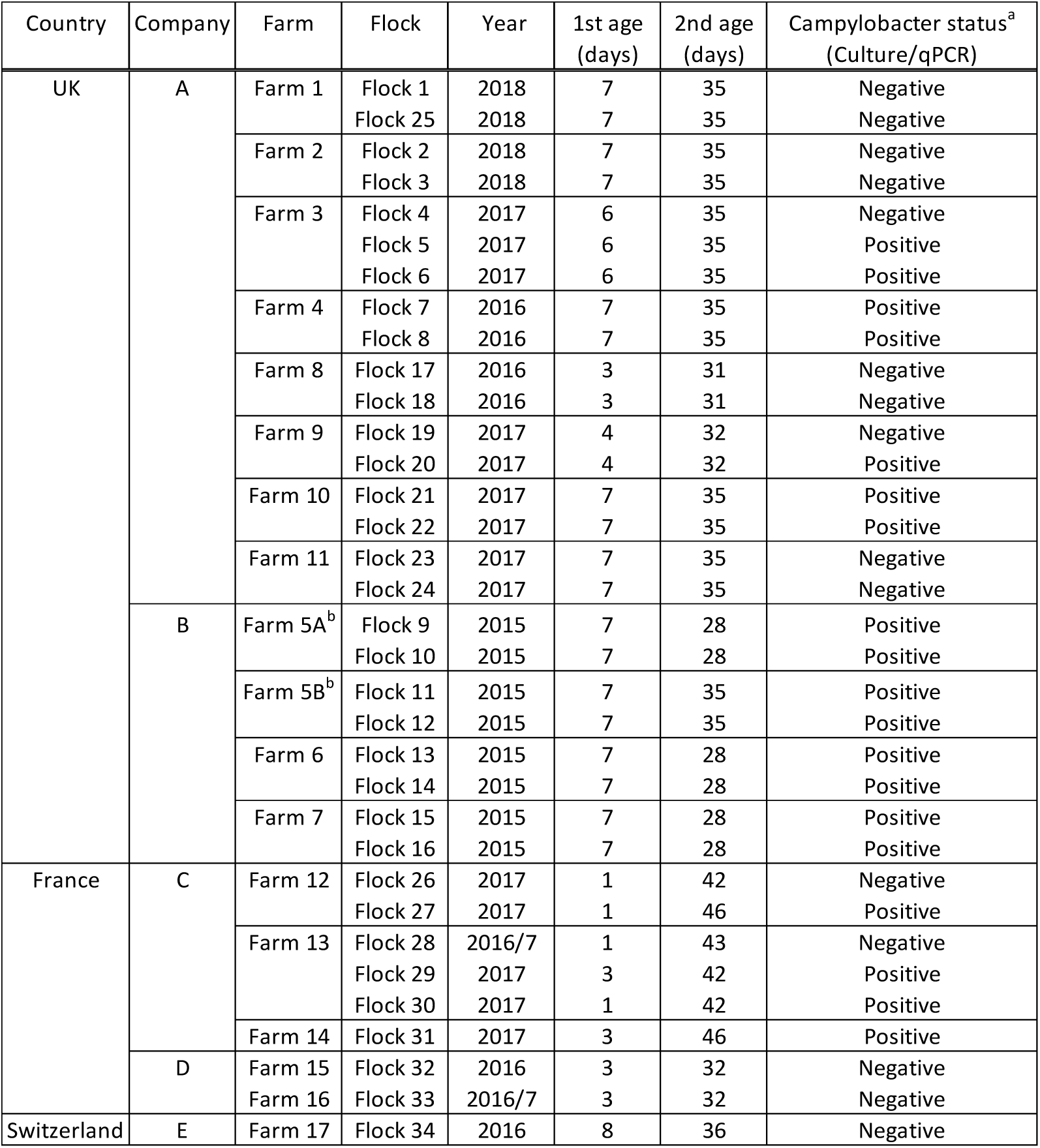
Details of the commercial broiler flocks from which faecal samples were tested. A total of 34 broiler flocks were tested from 3 countries and 5 companies, sampled between the years 2015-2018. Up to 5 fresh faecal samples were tested from different birds at early (< 8 days of age) and late (28-46 days of age) for each flock. ^a^Flocks were tested for *Campylobacter* by conventional culture or qPCR, using samples collected from the second sample date when birds were aged 28-46 days. ^b^ Flocks 11/12 were placed on Farm 5 after flocks 9/10 had left.

**Table 4.** The predominant *porAf2* types identified amongst the broiler flocks tested. The prevalence was compared between young birds from the same flock, with young birds from other flocks based upon total sequence data recovered per sample. *Predominant (>40%, most often >75%) of the total sequences recovered from a sample. Percent of birds was defined as the number of the birds from which the *porAf2* type was identified, divided by the number of young birds tested in a flock.

| Flock | Predominant *porAf2* type(s)*<br>(Percent of birds) | Average prevalence in young birds (1-7 days): |  |  |
| --- | --- | --- | --- | --- |
|  |  | Same flock<br>(Percent of birds) | Different flock: UK<br>(Percent of flocks) | Different flock: France<br>(Percent of flocks) |
| *UK flocks* |  |  |  |  |
| Flock 5 | 1 (75%) | 0.122 (100%) | 0.124 (100%) | 0.376 (100%) |
|  | 5 (50%) | 7.606 (100%) | 0.109 (96%) | 0.351 (100%) |
| Flock 6 | 1509 (100%) | 6.877 (100%) | <0.001 (75%) | Not detected |
| Flock 7 | 3 (100%) | 1.532 (100%) | 0.112 (100%) | 0.245 (100%) |
|  | 6 (33%) | 1.500 (100%) | 0.119 (100%) | 0.328 (100%) |
| Flock 8 | 3 (100%) | 2.209 (100%) | 0.111 (100%) | 0.245 (100%) |
| Flock 11 | 2170 (75%) | Not detected | <0.001 (8%) | <0.001 (14%) |
| Flock 12 | 2366 (33%) | Not detected | <0.001 (8%) | <0.001 (14%) |
|  | 687 (33%) | <0.001 (33%) | 0.003 (67%) | 0.001 (86%) |
|  | 2328 (33%) | Not detected | <0.001 (33%) | <0.001 (72%) |
| Flock 20 | 2342 (60%) | Not detected | <0.001 (13%) | <0.001 (14%) |
| Flock 21 | 7 (20%) | 3.59 (100%) | 0.190 (100%) | 0.581 (100%) |
|  | 2 (80%) | 1.407 (100%) | 0.040 (96%) | 0.638 (86%) |
| Flock 22 | 2 (80%) | 1.471 (100%) | 0.055 (96%) | 0.638 (86%) |
| *French flocks* |  |  |  |  |
| Flock 27 | 687 (100%) | 0.007 (100%) | 0.003 (67%) | 0.001 (86%) |
| Flock 29 | 2141 (60%) | 0.043 (80%) | 0.001 (38%) | 0.001 (86%) |
|  |  | 4.031 (80%) | 0.107 (96%) | 0.34 (100%) |
| Flock 30 | 8 (80%) | 6.64 (100%) | 0.048 (92%) | 0.127 (100%) |
|  | 3 (20%) | 1.479 (100%) | 0.108 (100%) | 0.228 (100%) |
| Flock 31 | 2141 (80%) | Not detected | 0.001 (38%) | 0.002 (86%) |
|  | 7 (20%) | 7.105 (100%) | 0.218 (100%) | 0.608 (100%) |

**Figure 2.**
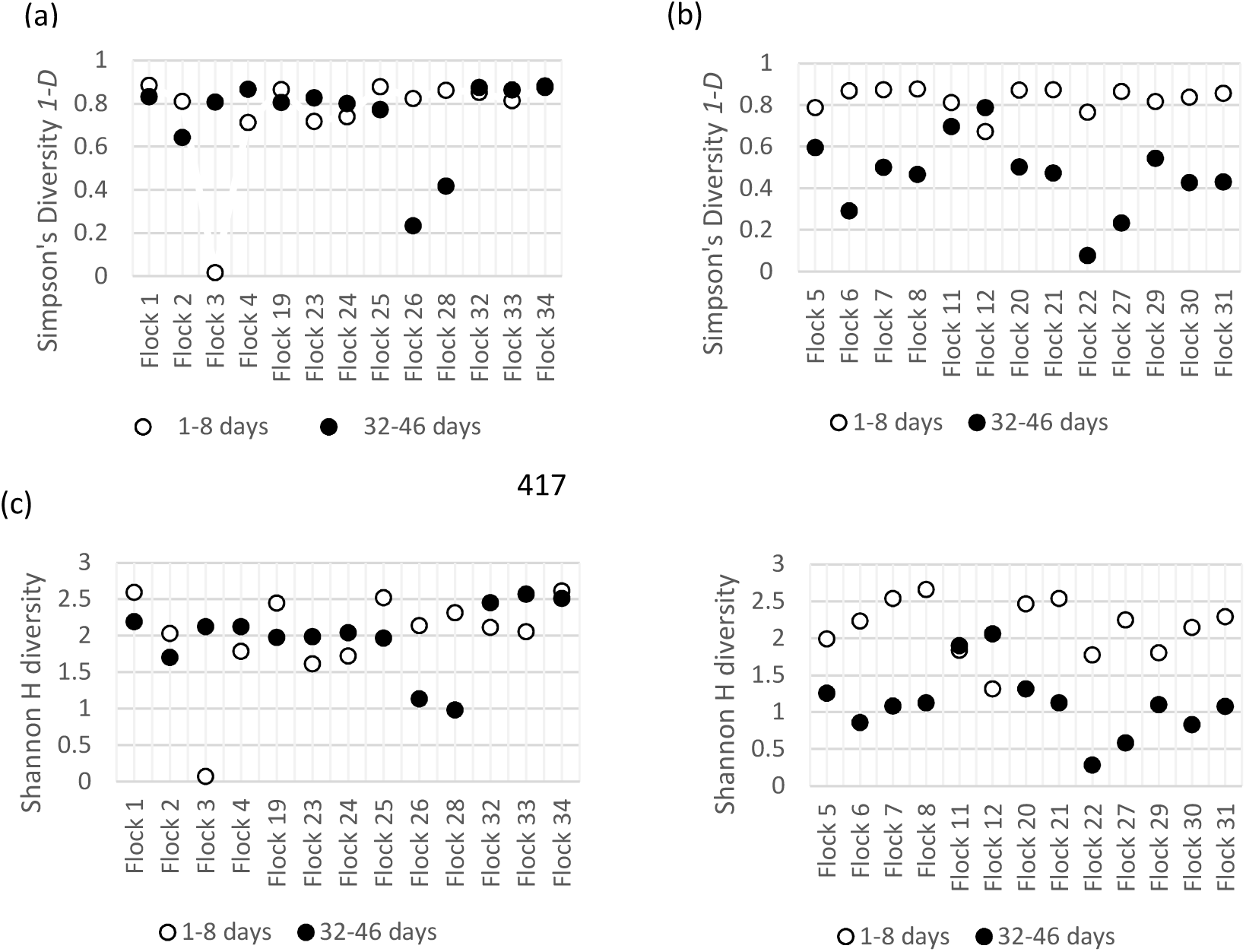
Diversity indices for *Campylobacter porAf2* variants identified amongst ‘*Campylobacter* negative’ flocks (a and c) and ‘*Campylobacter* positive’ flocks (c and d) using normalised data. Inverse Simpson’s diversity *1-D* (a) and (b) and Shannon H diversity (c) and (d).

Simpson’s and Shannon’s diversity indices were used to compare the *porAf2* genetic diversity of *Campylobacter* of flocks sampled at earlier and later time points (Figure 2). For both analyses, greater diversity was evident for samples from young birds, <8 days of age, compared to birds tested at 35-46 days of age from the same flock for *Campylobacter* culture positive flocks (Wilcoxon rank-sum exact test, N = 13 flocks, Simpson’s diversity *p* < 0.001, Shannon’s H diversity *p* = 0.002), but not for culture negative flocks (Wilcoxon rank-sum exact test, N = 13 flocks, Simpson’s diversity *p* 0.787, Shannon’s H diversity *p* = 0.68). No clustering of *porAf2* variant populations by country, company, farm or flock was evident on the Bray-Curtis ordination plots (supplementary data, Figure 6). Samples from older birds from ‘*Campylobacter positive*’ flocks had greater tendency to lie at the extremes of the plot, representing greatest differentiation of *porAf2* populations compared to other sample types.

When we compared methods, the *porAf2* results from 32-35 day old chickens showed that the *Campylobacter* detected by conventional culture matched those detected by deep sequencing (supplementary data, Figure 7). Of the 28 single-colony pick *Campylobacter* isolates cultured from birds aged 32-35 days, 18 (64.3%) matched the *porAf2* type that was predominant in the same sample by the deep sequencing method. For 9 samples, the predominant *porAf2* type detected by the deep sequencing method was not the same as the *porAf2* type detected by culture. Almost all of the cultured *porAf2* types could, however, be detected in the same sample by deep sequencing, at a percentage of the total *Campylobacter porAf2* types recovered, ranging from 0.12% to 46.99%. The one exception was *porAf2* type 2866 (Figure S5), which was cultured from flock 20/sample 5, but was not detected by deep sequencing from this sample.

## Discussion

The *porAf2* and 16S rRNA sequencing approaches show that *Campylobacter* genomic DNA is widely present in commercial broiler flocks in birds as young as 1-8 days of age, long before detection using standard methods. The very low abundance (0.0001-0.012%) of the microbiome DNA analysed provides an explanation for why these chicks are culture negative using standard methods. The early detection results are consistent with those from studies of cultured *Campylobacter* from 0.75% of 2,000 hatchery tray liners, representing pooled samples of 50 day of hatch chicks per tray liner^8^. Deep sequencing revealed great diversity of *porAf2* types in young birds, many of them present at less than 1% of the total. This level of resolution would require thousands of bacterial colonies to be sub-cultured to give even a small possibility of detection, which this is beyond the realistic capacity of standard culture methods. Although a wide range of genotypes were present in all samples; the proportions were more evenly distributed in young birds in low amounts. This contrasts markedly with older flocks that subsequently became *Campylobacter* culture-positive or PCR-positive from 28 days of age, where there was an order of magnitude more *Campylobacter* DNA in the sample with one or two dominant genotypes^5^. A striking result was that the *porAf2* analysis on the young bird samples showed no difference in either the abundance of *Campylobacter* present or in the range of genotypes between flocks that subsequently (from 28 days) became *Campylobacter* culture-positive and those that remained culture-negative. This suggests that all young flocks may have similar potential for growth of *Campylobacter* but that only some flocks experience the increase in bacterial growth that causes them to be *Campylobacter* culture or qPCR-positive.

The detection of *Campylobacter* in young birds alters the problem in the field from one of biosecurity during the growing phase, to the very youngest birds and the factors that affect *Campylobacter* outgrowth in some flocks. These findings suggest new lines of inquiry as to how broiler flocks become colonised by *Campylobacter* to the point where it becomes a threat to human health. First, it is important to determine how so many chicken flocks become contaminated with *Campylobacter* at this young age. Second, given the widespread presence of *Campylobacter* in young flocks, it is important to understand why some maintain low levels of *Campylobacter* throughout their lives and some exhibit the overgrowth of single types that result in *Campylobacter* culture-positive flocks. In addition to the biosecurity measures that have formed the poultry industry’s main line of defence against contamination, it may also be necessary to consider vertical sources of colonisation as well as the welfare and management of the birds themselves as key to controlling absolute levels of *Campylobacter* contamination in foods.

## Methods

### Faecal sample collection

Fresh faecal samples were collected in the period 2015-2018 from 34 commercially-grown broiler chicken flocks, representing: 17 farms; 5 companies; and 3 countries (Table 3). Samples from Company A (UK) were cultured for *Campylobacter* within 48 hours of collection, then stored at - 80°C. Samples from company B (UK) were frozen at −80°C within a day of collection and later tested by qPCR. Samples from companies C, D, and E (France and Switzerland) were immediately added to RNA*later*® (ThermoFisher Scientific, UK) and stored at −20°C. Faecal samples collected from ten SPF chickens at 3, 7, 14, 21, 35, and 49 days of age and stored at −80°C were employed as a negative controls.

### Detection of *Campylobacter* using culture or qPCR

Samples from company A were cultured on mCCDA (PO0119, Oxoid, UK) and incubated in a microaerobic atmosphere using the GenBox Microaer system (Biomérieux Ltd, UK) at 42°C for 48 hours. A single presumptive *Campylobacter* colony from each sample was sub-cultured onto Columbia Blood Agar (CBA) (PB0122, Oxoid, UK) and incubated for a further 48 hours at 42°C. Identity was confirmed by characteristic appearance and Gram stain, catalase and oxidase tests, and by DNA sequencing of the *porAf1* fragment (previously referred to as the *porA* fragment) using the Sanger sequencing method^5^.

Faecal samples from companies B, C, D and E were tested for the presence of *Campylobacter* using qPCR. Between 10 and 20 samples were pooled for each flock at each time point, and the DNA extracted using the protocols below. Primers and probes detecting portions of the *mapA* (*C. jejuni*) and *ceuE* (*C. coli*) genes, published by Best *et al*^9^, were used to detect the presence of *C. jejuni* and/or *C. coli.* Positive results were recorded for Ct values between 20 and 32.

### DNA extraction from faecal samples

Equipment and surfaces were decontaminated using 2% Virkon, DNA*Zap*TM PCR DNA degradation solutions (AM9890, ThermoFisher Scientific, UK) and 70% ethanol before use, and between batches. DNA was extracted from faecal samples not stored in RNA*later*® using the Maxwell 16 LEV Blood DNA kit (AS1290, Promega, UK) and automated DNA extraction system. Briefly, 250 mg of sample was added to a tube containing ¼ volume 0.5mm zirconia/silica beads (Thistle Scientific, UK) and 600µl lysis buffer. Samples were subject to a bead beater for 1 minute at high speed, and then heated at 95°C for 10 minutes in a water bath. They were centrifuged in a micro-centrifuge at 13,200 rpm for 5 minutes before transferring the supernatant to fresh tubes containing proteinase-K and incubating at 56°C for 20 minutes. The samples were then loaded onto the Maxwell 16 LEV Blood kit cartridges, and DNA extraction performed using the manufacturer’s ‘blood’ protocol.

Faecal samples stored in RNA*later*® were centrifuged at 13,200 rpm for 10 minutes and the supernatant removed prior to DNA extraction procedures. They were then washed in excess cold PBS and centrifuged for a further 10 minutes at 13,200 rpm and the supernatant discarded. DNA was then extracted using the Qiagen DNEasy PowerSoil kit (#12888-100, Qiagen, UK) following the manufacturer’s instructions. DNA prepared using the Maxwell system was diluted 1:10 before use to remove the effects of PCR inhibitors, but was used neat for Qiagen extractions

### Parallel sequencing for 16S (microbiome) and *porAf2* (*Campylobacter*)

Using a standard approach, 25µl PCR reactions were prepared in triplicate, within a PCR UV cabinet within a designated clean room. Equipment was decontaminated before use and between batches using RNAse Away (7002, ThermoFisher Scientific, UK), 70% ethanol and UV light for a minimum of 15 minutes. Fresh plastic ware and reagents/aliquots and non-template controls containing molecular water were used for every set of PCR reactions. 16S libraries were loaded onto a MiSeq sequencing run with the aim of getting 20,000 plus sequencing depth per sample (approximately double the depth of many studies), and *porAf2* libraries were loaded with the aim of getting 10,000 sequencing depth per sample.

For a subset of samples, variable regions 3 and 4 of the 16S ribosomal RNA was sequenced using the following primers: E338 5’-ACTCCTACGGGAGGCAGCAGT −3’ and R806 5’-GGACTACHVGGGTWTCTAAT −3’. For all samples, a short 405-473bp region of the *porA* gene was sequenced using the following primers: MOMP B 5’-CCA CAA TTA TGG TTA GCT TA −3’ and MOMP 2R 5’-TGA GAA GTT AAG TTT TGG AGA G-3’ ^5^. Either the forward or reverse primer was tagged with a different 7 or 10 nucleotide ‘barcode’ for each reaction, enabling them to be multiplexed within the same sequencing library. For both sets of PCR, the high fidelity Phusion Hot Start Flex DNA polymerase enzyme and 5X Phusion HF buffer (New England Biolabs, UK, M0535) were used according to the manufacturer’s recommendations. The following thermocycling conditions were used for; (1) 16S: initial denaturation 98°C for 5 minutes, 25 cycles of 98°C for 15 seconds, 50°C for 15 seconds, 72°C for 90 seconds, and final extension of 72 °C for 10 minutes and (2) *porAf2*: initial denaturation 98°C for 30 seconds, 35 cycles of 98°C for 10 seconds, 58°C for 30 seconds, 72°C for 30 seconds, and final extension of 72°C for 10 minutes. Library preparation was performed in a third room, separate to DNA extraction and PCR reaction setup, using protocols described previously^5^. PCR products were sequenced using the Illumina MiSeq platform, and 600-cycle MiSeq Reagent Kit v3 (Illumina, UK, MS-102-3003), giving paired 300 nucleotide reads. Samples were loaded with 10% phiX.

### Separately processed samples

A subset of 10 samples, from flocks 1 and 21 aged 7 days, were prepared individually through DNA extraction and PCR processes to control for potential cross-contamination between samples prepared in the same batch. At each stage, only one of the samples was handled in a room at a time and the process of DNA extraction was performed, or PCR completed for a particular sample, before the next sample was started. DNA extraction was performed for three samples from flock 1 using a Maxwell 16 robot (a single cartridge used at a time) and for two samples from flock 1, and five samples from flock 21 by using the QIAGEN DNEasy PowerSoil kit in enclosed tubes.

### Downstream processing of MiSeq data

Barcoded reads were demultiplexed and terminal primer sequences were trimmed using custom Python scripts. Any reads still containing a copy of either primer sequence after this trimming were identified using cutadapt v1.15, and excluded from further analyses on the assumption that they represent PCR artefacts (e.g. concatamers)^10^. The fastq_eestats2 command in USEARCH v10.0.240 was used to assess the optimal 3’ trimming length to remove low quality sequence, which was 270bp for our data ^11^. The DADA2 3.10 software package was used to trim reads, filter to <=1 expected errors/read, assign sequence variants (using pooled sequences), merge read-pairs, and remove chimeras (using the removeBimeraDenovo() function with method = “consensus”)^12^. The count of *Campylobacter* 16S OTUs per sample were calculated from a Phyloseq object using the subset_taxa function^13^. The *porAf2* nucleotide sequences were aligned in MEGA-X to confirm that they were coding sequence for correct fragment from *Campylobacter*^14^. A small number of sequences containing stop codons were removed from further analyses. Allele numbers for nucleotide and peptide sequences for the *porAf2* were assigned using the PubMLST *Campylobacter* database (https://pubmlst.org/campylobacter)^15^.

### Data analysis

Data were normalised to compare the diversity of *Campylobacter porAf2* types amongst samples. This was achieved by transforming the total number of counts per sample to relative abundance using the transform_sample_counts function in R^16^. Data from Flock 14, aged 7 days, were removed from the analyses at this point as the sequencing depth was between 127 and 281. Bar charts and Table 3 were prepared using Tableau 2019.4 software. Paired Wilcoxon Rank-Sum Exact tests were used to compare (i) the difference in percent microbiome represented by *Campylobacter* 16S rRNA between flocks tested at young and older time points; (ii) the prevalence of a *porAf2* type amongst young birds in a flock before it became predominant, with the average prevalence amongst young birds from other flocks; (iii) the number of *porAf2* types identified amongst flocks tested at young and older times points; and (iv) Simpson’s diversity index and Shannon’s H diversity index between *Campylobacter* culture positive and negative flocks. A Mann-Whitney U test was used to compare the number of *porAf2* types identified amongst *Campylobacter* culture positive and negative flocks. The tests were performed using Paleontological statistics (PAST) version 4.0 software^17^.

PAST version 4.0 software was also used to calculate a modified Simpson’s diversity index (1-*D*) and Shannon’s H diversity index (normalised data) and rarefaction curves for samples pooled by flock/age (total sequencing data). For Simpson’s diversity index, a 1-*D* value of 1.0 indicated that all members of a population could be distinguished from each other, and a 1-*D* value of 0 indicated that all members of a population were identical^18^. Shannon’s diversity index was included as it is considered to give more weight to rare species (*pofAf2* variants)^19^. An H value of 0 indicated that all species were the same. H increases with increasing number of species.

The Bray-Curtis dissimilarity index was used to compare the populations *of Campylobacter porAf2* genotypes between birds aged 1-7 days, and 31-46 days from different UK and French flocks^20^. A Bray-Curtis value of 0 meant that two flocks have the same *Campylobacter porAf2* population and a value of 1 meant that two flocks did not share any *porAf2* types. Data were normalised using the transform sample counts function in R as above and then converted into a Phyloseq object^13^. The ordination function with NMDS method was used in R to produce the Bray-Curtis ordination plots.

### Ethical approval

This study used broiler flocks that were reared commercially by industry partners, in line with standard industry practice and involved non-invasive sampling of faeces. All prevailing local, national and international regulations and conventions, and normal scientific ethical practices have been respected. The Swiss samples were approved by the Canton of Bern (BE97/16) and met all cantonal and federal regulation for the treatment of animals.

## Acknowledgements

This work was supported by the Biotechnology and Biological Science Research Council (grant numbers BB/N023803/1 and BB/K004468/1) including part of the Animal Health and Welfare ERA-net call, and also the BBSRC DTP Integrated Biosciences (grant number BB/M011224/1); the United Kingdom Department for Environment, Food and Rural Affairs (grant number OD0221), the United Kingdom Food Standards Agency [grant number FS101013]; the Wellcome Trust [grant number 087622]; the National Institute for Health Research Health Protection Research Unit (NIHR HPRU) in Gastrointestinal Infections at the University of Oxford in partnership with Public Health England (PHE) and the Bertarelli Foundation. The views expressed are those of the author(s) and not necessarily those of the BBSRC, FSA, NHS, the NIHR, The Bertarelli foundation, Department of Health or Public Health England. We gratefully acknowledge in-kind support from Avara Foods, and thank all companies for providing access for samples. We also thank Prof. Dr. Benjamin Schusser from Technical University of Munich, who may be an author in future versions of this manuscript, for allowing us to test faecal samples from SPF birds.

