## Supplementary data for "Deep sequencing reveals *Campylobacter* in commercial meat chickens less than 8 days old"

**Figure S1. The total number of 16S rRNA sequences obtained (coverage) and count of *Campylobacter* 16S sequences (*Campylobacter* count) from a subset of 16 UK broiler flocks.** Data are broken down by (a) flocks, and (b) individual birds. *Campylobacter* counts up to a value of 10 are shown for purposes of scale. Less than 10 *Campylobacter* 16S sequences were recovered from birds aged 6-7 days. Eleven samples from birds aged 28-35; Flock 5, bird 6 (n=14), bird 8 (n=584); Flock 6, bird 6 (n=45), bird 10 (n=16); Flock 8, bird 7 (n=35), bird 8 (n=20); Flock 11, bird 8 (n=14); Flock 12, bird 6 (n=87), bird 9 (n = 21); Flock 15, bird 10 (n=16) and Flock 16, bird 8 (n=59).

**(a)**

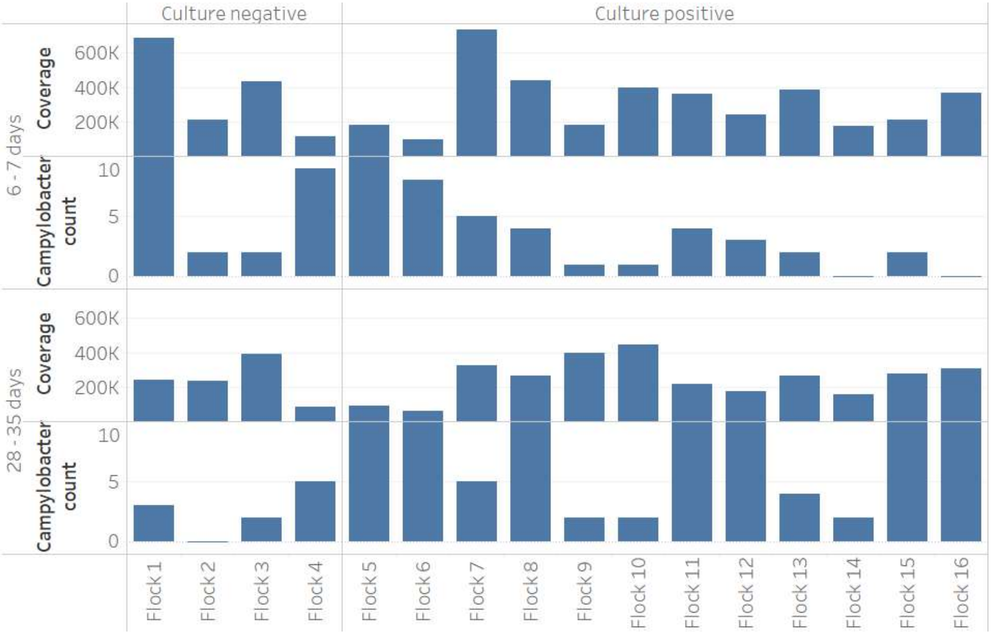


**(b)**

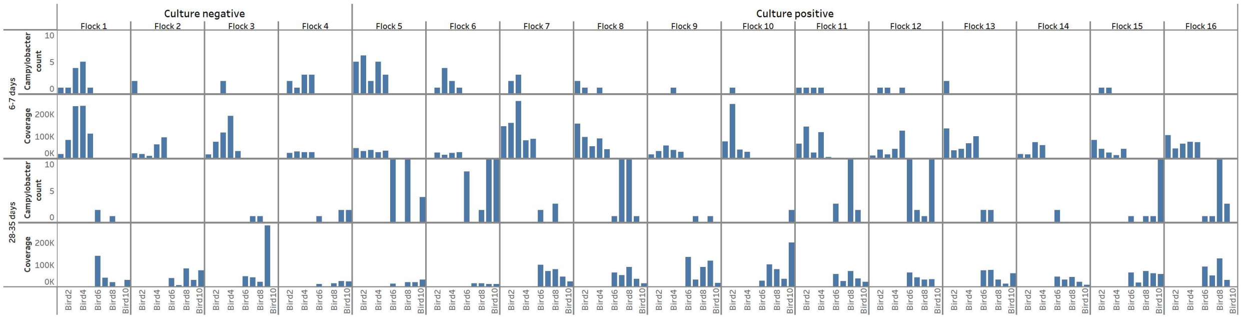

**Figure S2. Rarefaction curves for *Campylobacter porAf2* types isolated from up to 5 pooled faecal samples/flock for commercial broiler flocks from the UK, France and Switzerland.** Results are grouped by bird age; (a) young birds 1-8 days and (b) older birds 28-46 days.

(a)

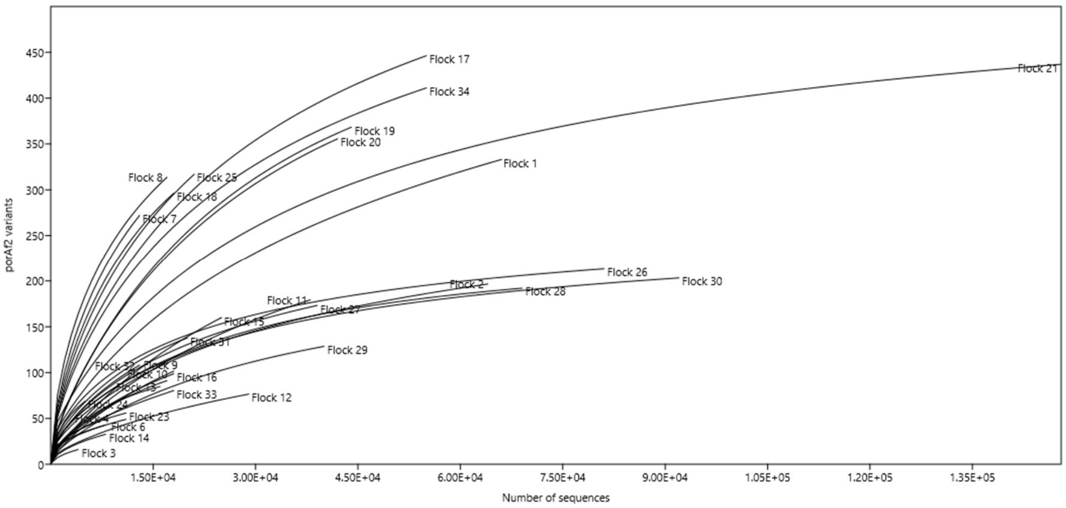


(b)

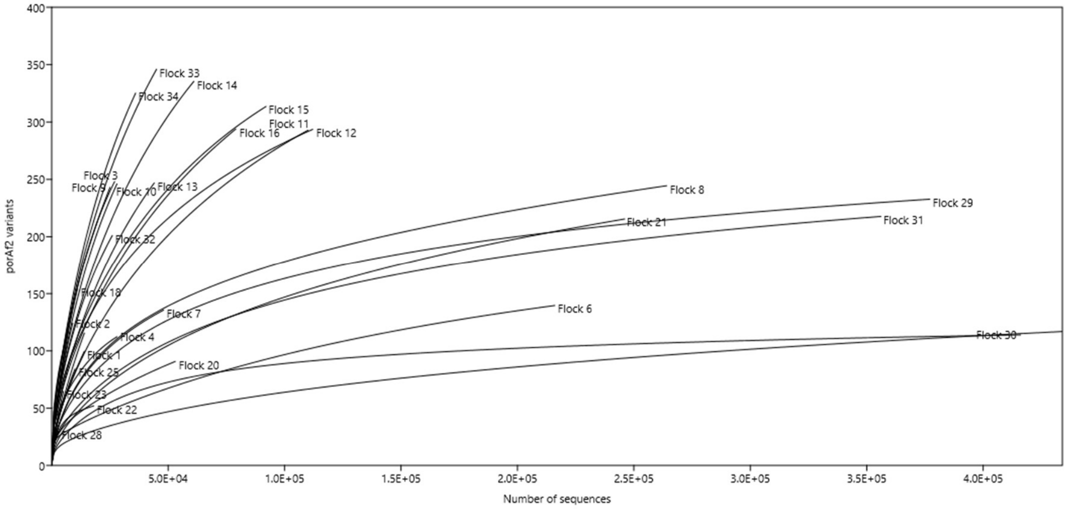

**Figure S3. The cumulative total of *Campylobacter porAf2* variants identified per flock.** The asymptotic curve indicates that sampling captures most of the variants across the three countries.

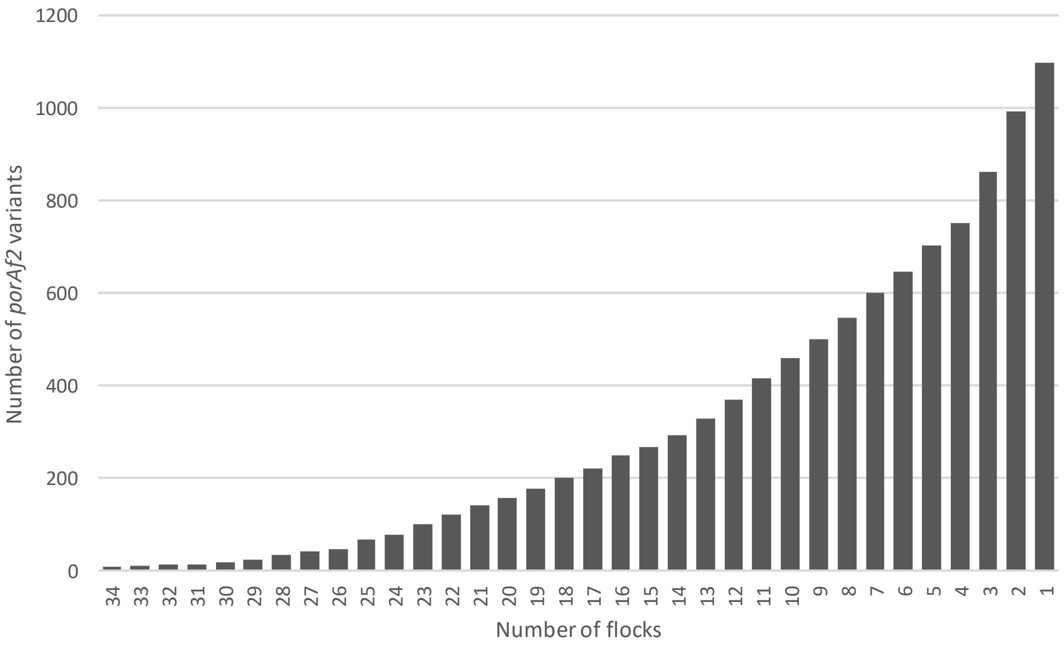

**Figure S4. A neighbour joining tree showing evolutionary distance amongst 1098 *Campylobacter porAf2* variants isolated from commercial broiler flocks.** The most commonly isolated *porAf2* variants (1-7) are highlighted in colour.

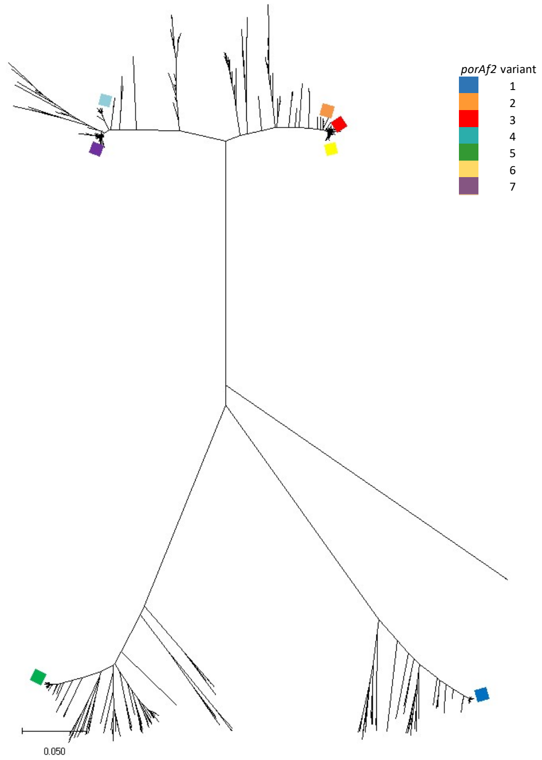

**Figure S5 The number of *Campylobacter porAf2* variants recovered relative to sequencing depth for (a) *Campylobacter* culture negative *vs* culture positive flocks and (b) compared to flock age (days) (normalised data).**

(a)

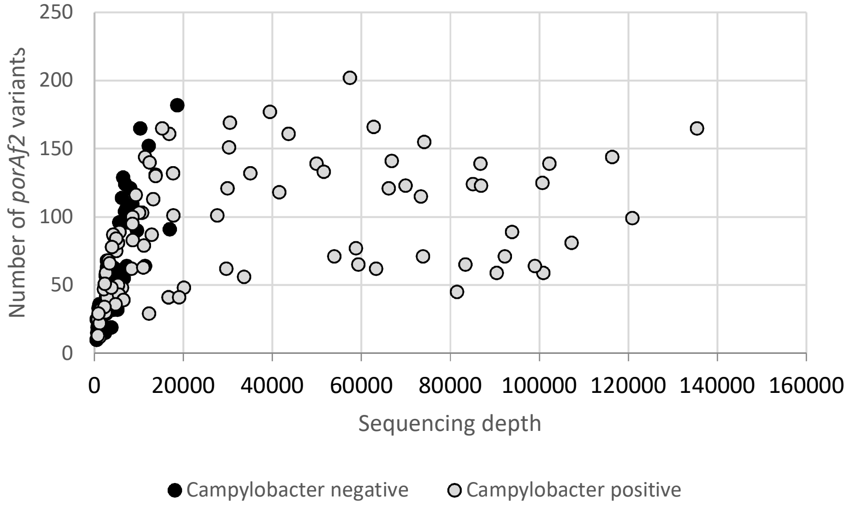


(b)

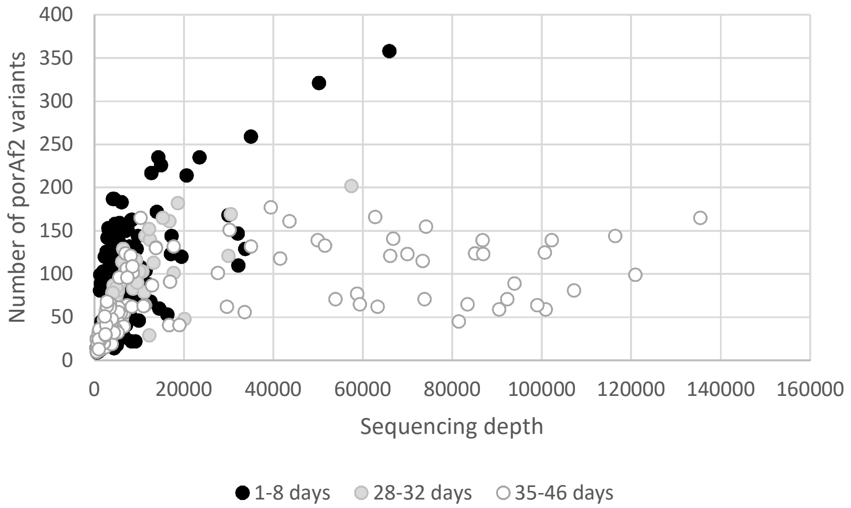

**Figure S6. Ordination plots showing Bray Curtis dissimilarity indices for *Campylobacter porAf2* variants from commercial broiler flocks, separated by *Campylobacter* status (culture/qPCR) of the sample, Country, Company, Farm, Flock and Age.**

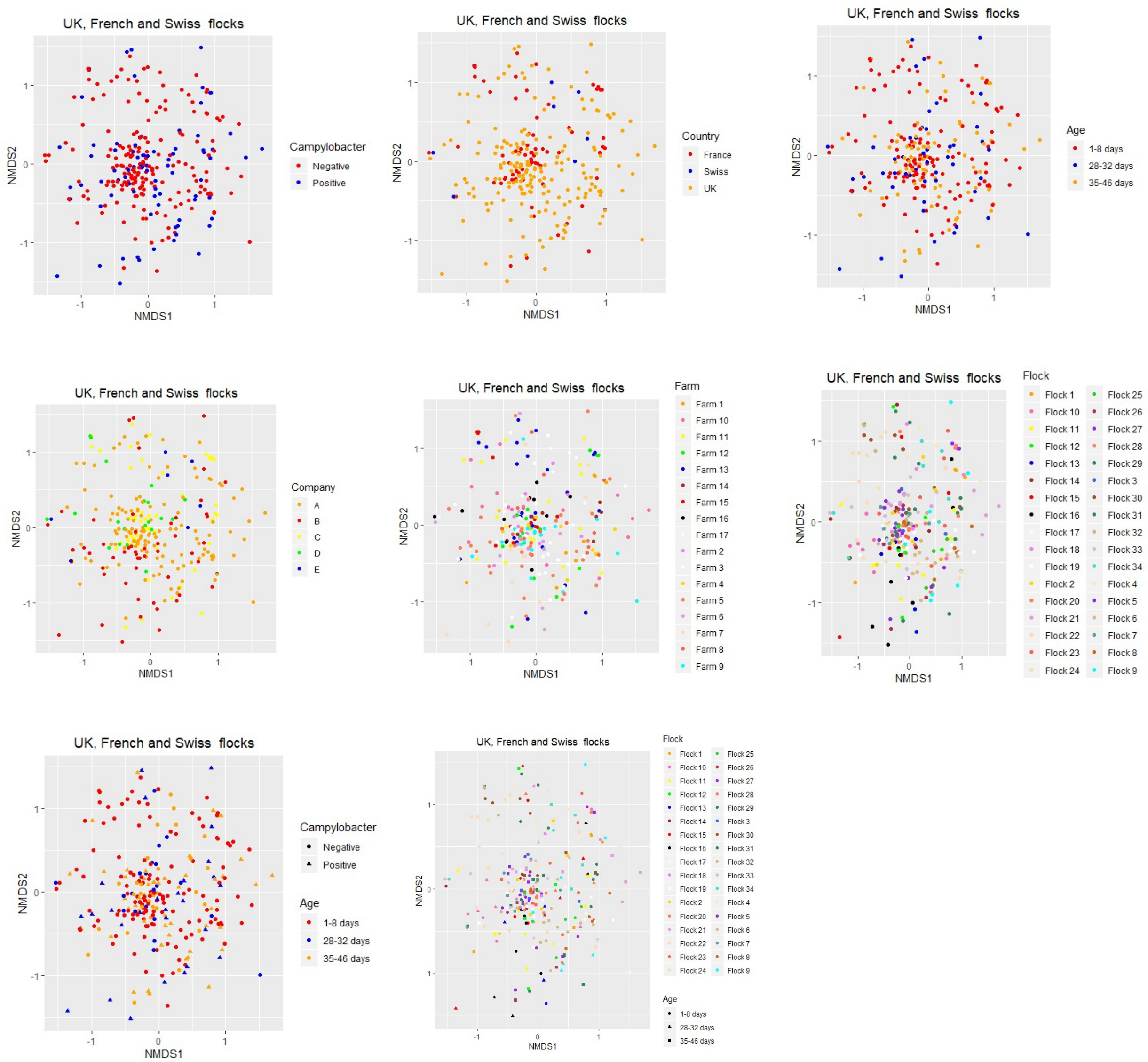

**Figure S7. Comparison of *Campylobacter porAf2* types identified amongst samples using (a) deep sequencing and (b) conventional culture amongst a subset of flocks.** Each colour represents a different *porAf2* type; common variants are given in the legend.

(a)

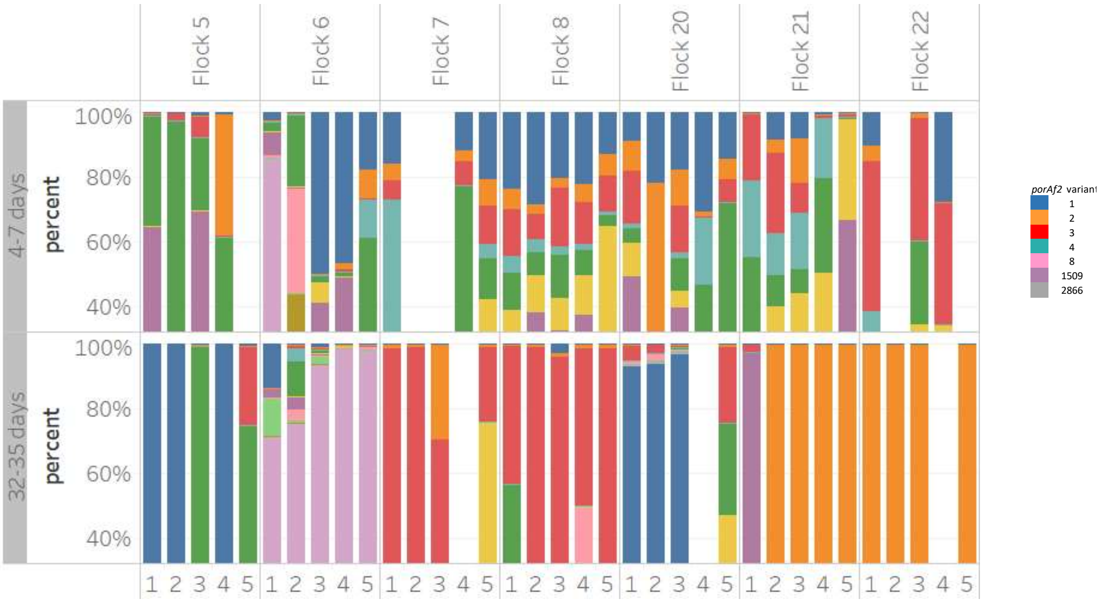


(b)

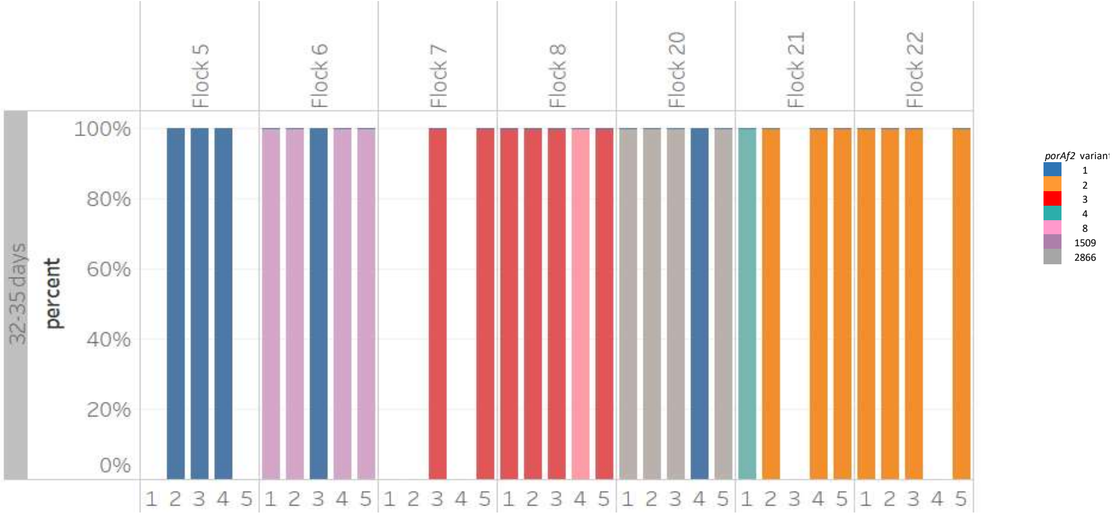
